# Spatiotemporal cortical dynamics for rapid scene recognition as revealed by EEG decoding

**DOI:** 10.1101/2023.02.16.528781

**Authors:** Taiki Orima, Isamu Motoyoshi

## Abstract

The human visual system rapidly recognizes the categories and global properties of complex natural scenes. The present study investigated the spatiotemporal dynamics of neural signals involved in ultra-rapid scene recognition using electroencephalography (EEG) decoding. We recorded visual evoked potentials from 11 human observers for 232 natural scenes, each of which belonged to one of 13 natural scene categories (e.g., a bedroom or open country) and had three global properties (naturalness, openness, and roughness). We trained a deep convolutional classification model of the natural scene categories and global properties using EEGNet. Having confirmed that the model successfully classified natural scene categories and the three global properties, we applied Grad-CAM to the EEGNet model to visualize the EEG channels and time points that contributed to the classification. The analysis showed that EEG signals in the occipital lobes at short latencies (approximately 80∼ ms) contributed to the classifications other than roughness, whereas those in the frontal lobes at relatively long latencies (∼ 164 ms) contributed to the classification of naturalness and the individual scene category. These results suggest that different global properties are encoded in different cortical areas and with different timings, and that the encoding of scene categories shifts from the occipital to the frontal lobe over time.

## Introduction

It is widely known that the human visual system rapidly discriminates complex natural scenes and utilizes perceived information to judge the surrounding environment and for spatial navigation (Potter, 1975; Intraub, 1981; Thorpe et al., 1996; Oliva & Torralba, 2001; Greene & Oliva, 2009; Peelen et al., 2009; Bonner & Epstein, 2018). This rapid perception is thought to be based on information obtained by a glance at a natural scene, which precedes the perception of individual objects or detailed features within the scene. Such information is often referred to as gist, which has been successfully formulated as a relatively global image feature such as the spatial envelope (Oliva & Torralba, 2001; Torralba & Oliva, 2003). Specifically, the spatial envelope is a low-order feature designed to provide a good estimate of the degrees of important indicators with which to characterize a class of natural scenes, such as naturalness, openness, roughness, expansion, and ruggedness (Oliva & Torralba, 2001). Some of these indicators can be discriminated with high accuracy even when the visual stimuli are presented for 50 ms or less (Joubert et al., 2007; Greene & Oliva, 2009). However, these behavioral data have various factors beyond visual processing of the target scene itself, such as the properties of the backward masking effect (Breitmeyer et al., 2006) and the decision process for response selection (Shadlen & Kinani, 2013; Ratcliff et al., 2016). Analyzing the brain activities for natural scene images may enable us to understand the dynamics of scene processing in humans more directly.

Neural mechanisms of scene perception in the human brain have been most extensively investigated through functional magnetic resonance imaging (fMRI). Comparisons of blood oxygenation level dependent signals between visual stimuli having specific characteristics, such as scenes and faces, have revealed scene-selective regions that are important for the perception of natural scenes. The parahippocampal place area is located from the posterior part of the parahippocampal gyrus to the anterior part of the spindle gyrus and has been identified as a region that shows preference for buildings (e.g., Epstein & Kanwisher, 1998; Aguirre et al., 1999). The retrosplenial complex, which is active against mental images of the scene, and the occipital place area, which shows preference for the boundaries of the environment in navigation, have also been identified as scene-selective regions (O’Craven & Kanwisher, 2000; Nakamura et al., 2003; Julian et al., 2016; Groen et al., 2016; Bonner & Epstein, 2018; Epstein & Baker, 2019). These findings suggest that multiple areas in the human brain process different types of information from natural scene images. However, because of the low temporal resolution of fMRI, the cited work could not specify the early neural activities corresponding to rapid natural scene processing, which is probably based on image features as suggested by a number of psychophysical and computational studies (Schyns & Oliva, 1994; Baddeley, 1997; Oliva et al., 1999; Oliva & Torralba, 2001; Gasper & Rousselet, 2009).

Meanwhile, the temporal dynamics of neural processing underlying rapid scene recognition have been investigated through electroencephalography (EEG). A recent study showed that differences in the global information of natural scenes evoked different visual evoked potentials (VEPs) (Harel et al., 2016; Hansen et al., 2018). Another line of research has focused on the hierarchical neural processing of image features that are important for scene recognition. Focusing on a lower-order feature called contrast energy and a higher-order feature called the spatial coherence of natural scene images, Groen et al. (2013) showed that the modulation of EEG by contrast energy terminated in 100–150 ms, whereas the modulation by spatial coherence lasted up to 250 ms. Greene & Hansen (2020) investigated the relationship between event related potentials (ERPs) and a wide range of features from lower to higher order (i.e., features ranging from simple texture statistics of natural scenes to convolutional neural network (CNN) features) and found differences in the encoding process for each feature. Referring to a large body of evidence suggesting that the important features for the instantaneous perception of natural scenes are relatively global (Oliva & Torralba, 2001; Greene & Oliva, 2009; Groen et al., 2013; Kauffmann et al., 2014; Ramkumar et al., 2016), it has been suggested that the natural scene encoding process at an early stage can be investigated using EEG (Scholte et al., 2009; Ghebreab et al., 2009; Võ & Wolfe, 2013; Groen et al., 2016). However, the low spatial resolution of EEG does not allow us to investigate the spatial distribution of the scene-related neural activity over the cortex.

Although various psychophysical and neurophysiological approaches have been adopted to examine the perception of natural scenes, it remains unclear, both spatially and temporally, what part of the brain activity at short latencies contributes to the classification of natural scene categories and global properties. In the present study, we conducted experiments to investigate the spatiotemporal development of neural information related to scene categories (e.g., a bedroom and forest) and fundamental global properties (i.e., the degrees of naturalness, openness, and roughness) using VEPs. We trained the EEGNet model (Lawhern et al., 2018), which was a CNN model that predicted the natural scene categories and global properties (degrees of naturalness, openness, and roughness) of corresponding natural scene images to inputting VEPs, and visualized the VEP time points and EEG channels that contributed to the classification using Grad-CAM (Selvaraju et al., 2017). These analyses showed that the corresponding natural scene categories and global properties could be classified from simple VEPs at a statistically significant level, and they visually revealed that the different time points and EEG channels contributed to different classification classes. In particular, we found that early-latency (approximately 80∼ ms) VEPs contributed to the naturalness and openness classifications, and that both frontal and occipital electrodes contributed to the naturalness classification. These results suggest that different global properties, which have been considered to be important for natural scene recognition, are processed in different cortical areas, and that their localization has already occurred within a short latency of ∼ 100 ms. In addition, these findings further support the idea that EEG with poor spatial resolution can be adopted to explore the dynamic neural processing of complex natural images.

## Experiment

We measured VEPs for various natural scene images and constructed an EEGNet model using the VEPs as input. We examined how accurately the model classified the natural scene categories and global properties of corresponding images. We then applied Grad-CAM to the EEGNet models to visualize the time points and EEG channels of the VEPs that contributed to the classification.

## Methods

### Observers

Twelve naïve students participated in the experiment. All participants had normal or corrected-to-normal vision. All experiments were conducted in accordance with the guidelines of the Ethics Committee for experiments on humans at the Graduate School of Arts and Sciences, The University of Tokyo. All experiments were conducted in accordance with the Declaration of Helsinki. All participants provided written informed consent. One participant was excluded from the following analyses because their EEG data were deficient.

### Apparatus

Visual stimuli were generated by a personal computer (HP Z2 Mini G4 Workstation) and presented on a 24-inch gamma-corrected liquid-crystal display (BenQ XL2420T) with a refresh rate of 60 Hz and a spatial resolution of 1.34 min/pixel at a viewing distance of 100 cm.

### Stimuli

The visual stimuli were 232 natural scene images, which subtended 5.7 deg x 5.7 deg (256 x 256 pixels) (Figure 1). All images were collected via the Internet from the SUN and Places 365 databases (Xiao et al., 2010; Zhou et al., 2014). All images were classified into one of 13 natural scene categories identified as important in previous studies: offices, kitchens, living rooms, bedrooms, industrial scenes, tall buildings, city scenes, streets, highways, coasts, open country, mountains, and forests (Oliva & Torralba, 2001; Lazebnik et al., 2006; Alameer et al., 2016).

**Figure 1.**
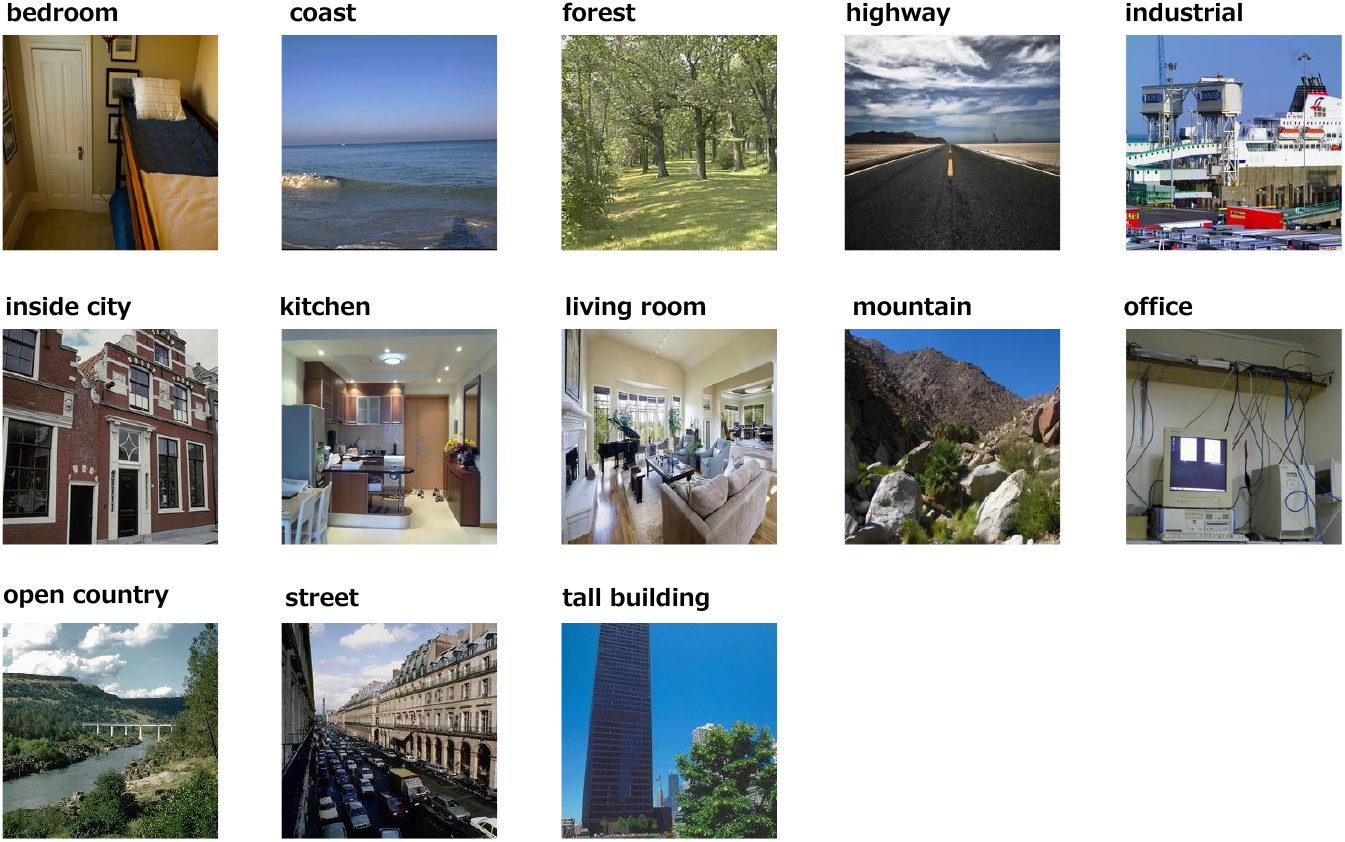
Examples of stimuli used in the experiment; i.e., one image from each of the 13 natural scene categories.

### EEG recording procedures

EEG experiments were conducted in a shielded dark room. In each session, 232 natural scene images were presented once in random order. Each image was presented for 500 ms, after which a uniform background blank was presented for approximately 750 ms. Participants observed the stimuli foveally through steady fixation on a small black dot that appeared at the center of the screen. EEG recordings were made while the participants observed the visual stimuli. Participants’ eye movements were controlled by pre-experiment instruction (c.f., Orima & Motoyoshi, 2021). 17 sessions were conducted in the experiment, and each image was presented 17 times in total for each participant.

### EEG data preprocessing

EEG data were acquired from 19 electrodes (Fp1, Fp2, F3, F4, C3, C4, P3, P4, O1, O2, F7, F8, T7, T8, P7, P8, Fz, Cz, and Pz) in accordance with the international 10-20 system at a sampling rate of 1000 Hz (BrainVision Recorder, BrainAmp amplifier, EasyCap; Brain Products GmbH). The impedance of each electrode was kept below 5 kΩ. An additional electrode, located between Fz and AFz, was used as the ground. In addition, all electrodes were referenced to another electrode located between Fz and Cz, and all electrode data were re-referenced offline using the average of all electrodes. The recorded EEG data were filtered by a 0.5–40 Hz bandpass filter and divided into epochs of -0.4-0.8 sec from the stimulus onset. Baseline correction was performed using the data for -100-0 msec from the stimulus onset as a baseline. The artifact components (i.e., eye movements) were removed through independent component analysis.

### Training the EEGNet model

EEGNet is a CNN model that treats EEG data as two-dimensional data of time points x EEG channels as input (Lawhern et al., 2018; Lotte et al., 2018). Previous studies have shown that EEGNet performs well in EEG decoding, and because it convolves both in time and in space, it is said to be able to capture the spatiotemporal properties of EEG data.

Grad-CAM has been used to visualize the portion of inputs that contribute to classification in deep neural network models for object recognition (Selvaraju et al., 2017). In the present study, not only to classify the characteristics of visual stimuli from VEPs but to understand the spatiotemporal portions that contributed to the classification, we trained an EEGNet model to classify corresponding natural scene categories and global properties, and applied Grad-CAM to the EEGNet model to visualize the classification.

Figure 2 is an overview of the EEGNet model. Following a previous study (Lawhern et al., 2018), EEG data were input as two-dimensional data of time points x EEG channels and trained to classify 13 natural scene categories of the corresponding visual stimuli to the input VEPs. The 232 images were split into training and testing data such that they were almost equally divided within each natural scene category. Five validation sets were prepared, and we performed five times cross validation.

**Figure 2.**
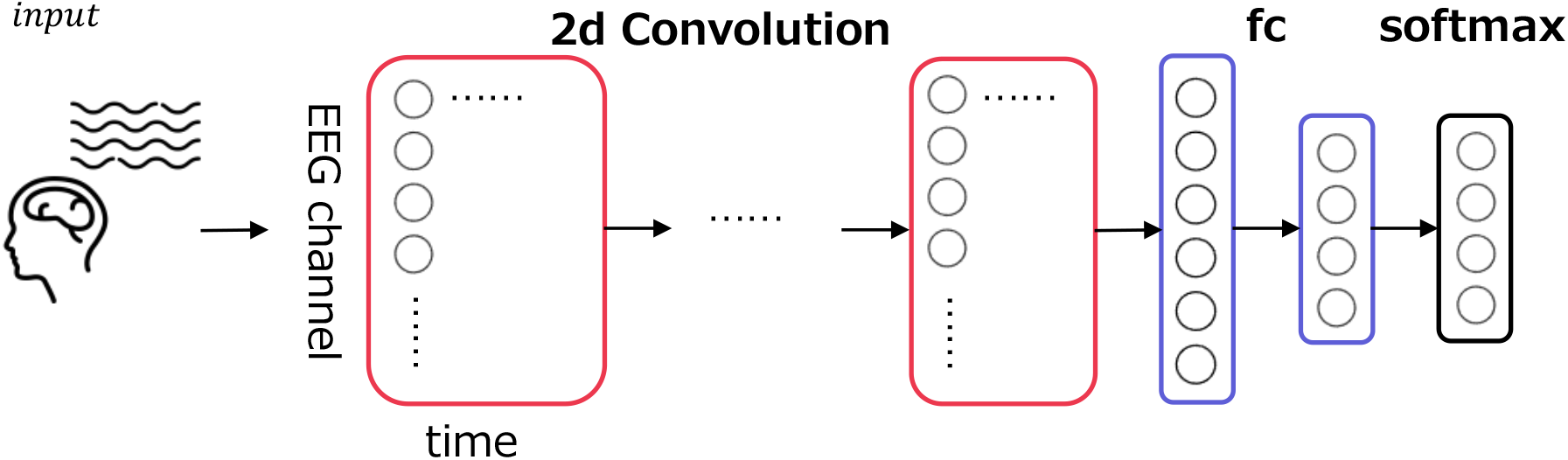
Model overview. EEG data were input as a two-dimensional array comprising 17 EEG channels x 500 time points to the convolutional layers. The convolutional layers were followed by a fully connected layer and then a softmax layer to classify natural scene categories and global properties.

The preprocessed EEG data from 1 to 500 ms of the stimulus onset for 17 electrodes (F3, F4, C3, C4, P3, P4, O1, O2, F7, F8, T7, T8, P7, P8, Fz, Cz, and Pz) in the international 10-20 system were treated as 17 x 500 matrix data. From ∼ 187 samples (11 observers x 17 repetition, some of them were rejected by the preprocess), thirty to thirty-five samples of EEG data corresponding to a single visual stimulus were picked up in random combinations for input to the model and then averaged. Note that each sample for each image, EEG channel and repetition were z-scored to eliminate the effect of the absolute value of each channel. The number of training epochs was set at 1,000, and the number of samples used for training per epoch was 3,000. Table 1 shows the detailed architecture of the EEGNet model. The loss for each iteration was calculated using PyTorch’s torch.nn.CrossEntropyLoss.

**Table 1.**
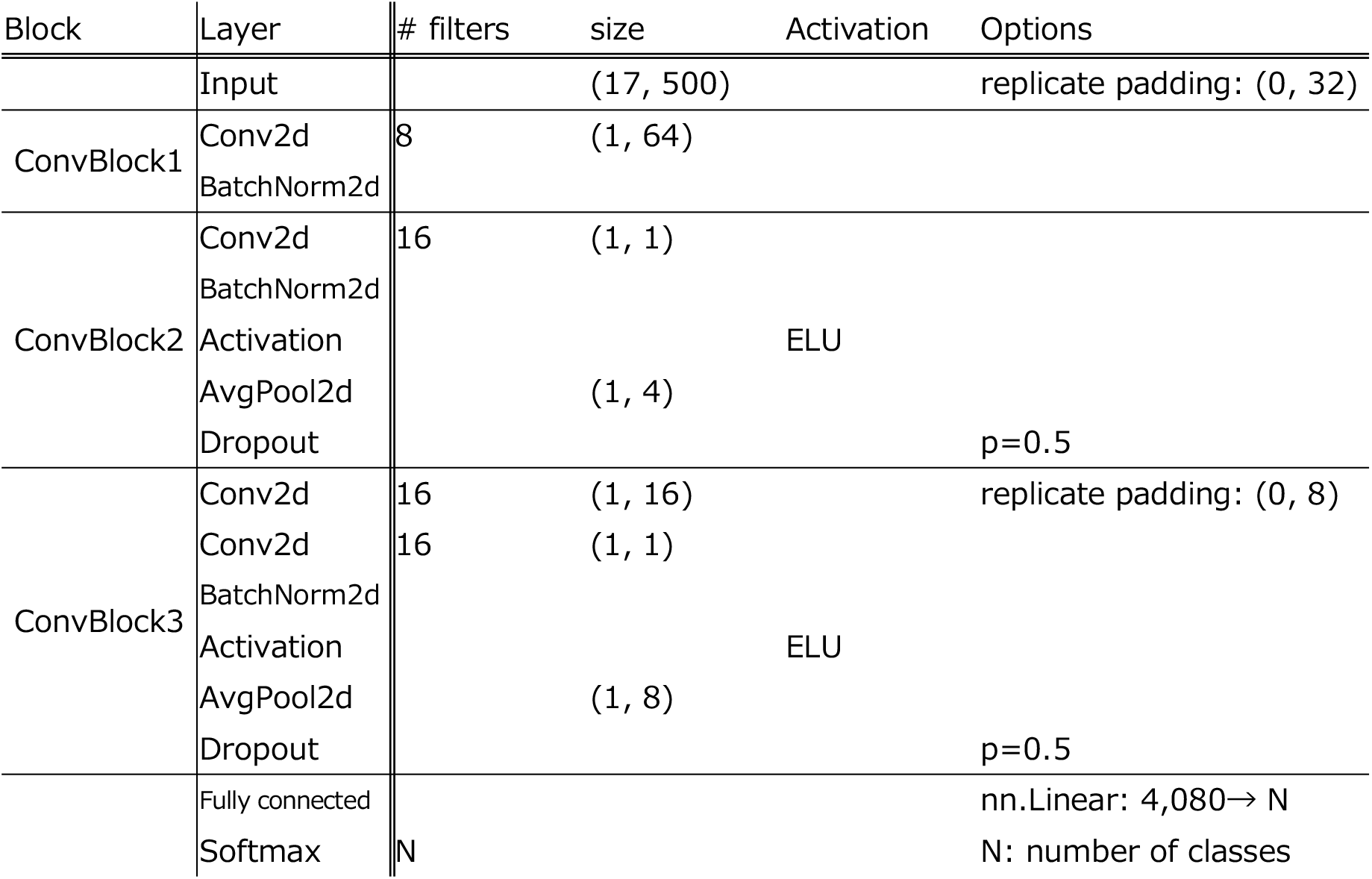
Details of the EEGNet architecture

Besides training the EEGNet model to classify the VEPs into corresopnding 13 natural scene categories, we also trained the EEGNet model to classify the VEPs according to global properties that characterize natural scenes, namely the degrees of naturalness, openness, and roughness (Oliva & Torralba. 2001). Naturalness (natural/man-made) had a predefined 0/1 value indicating whether each natural scene was mainly composed of natural or man-made objects. Openness (open/closed) also had a predetermined 0/1 value, indicating whether each natural scene was open or closed. Roughness (simple/complex) was considered to correspond to the ‘complexity’ of the scene (Oliva & Torralba, 2001). In the present study, the slope of the power spectrum of each image, which is related to roughness, was calculated and binarized around its median value to give the roughness of each image (Oliva & Torralba, 2001). The degrees of expansion and ruggedness were excluded from the present study because they are mainly applied only to man-made and natural scenes, respectively.

The architecture of the EEGNet models is the same as that shown in Table 1, except for the size of the final softmax layer. The number of training epochs was set at 1000, and the number of samples used for training in one epoch was set at 3000.

After the training of the EEGNet models, Grad-CAM was adopted to visualize the contribution to the classification. The average VEPs of each participant′s testing data were input to the trained EEGNet model, and the predicted natural scene categories were obtained from each VEP. We then applied Grad-CAM to the predicted indices following a previous study (Selvaraju et al., 2017). The output of the convolutional layer in ConvBlock2 was used as the feature map. Next, error back propagation was applied to the indices of the predicted natural scene category or global property, and the global average pooling of the feature maps was calculated. A localization map was obtained as the multiplicative product of the feature maps and global average pooling. To adopt only the points that contributed positively to the classification, the localization map was finally passed through a ReLU layer. The localization maps that were obtained for each participant were averaged across participants and projected onto a topographical map to visualize the time points and EEG channels that contributed to the classification.

## Results

### VEPs

Figure 3 shows the grand-average VEPs for all images from 50 to 500 ms after the stimulus onset. Red indicates positive amplitudes and blue indicates negative amplitudes. VEPs were particularly large for the occipital electrodes (O1, O2). VEPs of the occipital electrodes (O1, O2) began to rise at approximately 100 ms after the stimulus onset. The amplitudes of the VEPs of the occipital electrodes increased again, peaked at approximately 250 ms, and then decreased.

**Figure 3.**
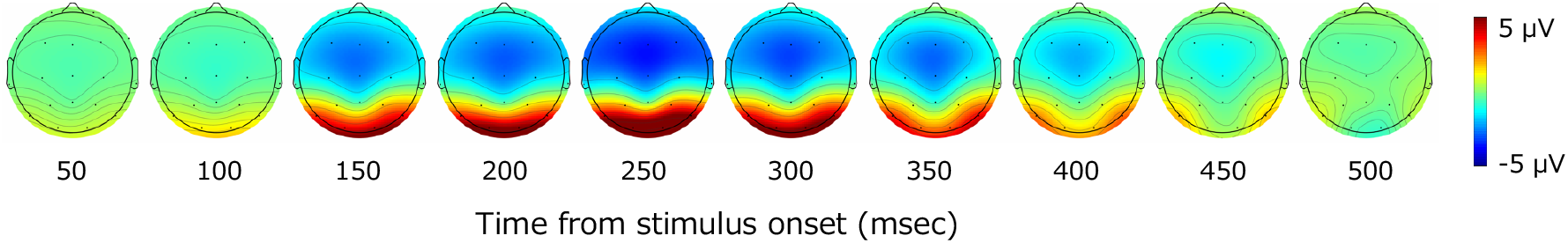
Topography of grand-average VEPs. Red indicates positive values and blue indicates negative values. A large rise in VEPs was observed after 100 ms from the stimulus onset, mainly in the occipital cortex.

### Classification of natural scene categories and global properties using EEGNet models

Figure 4 shows the classification accuracy d’ of the scene categories using the EEGNet model. In each cross-validation set, the VEPs that were assigned to the testing data were averaged within participants and input to the trained EEGNet model, and we obtained the classification accuracy for each participant and cross-validation set. The obtained values of classification accuracy were averaged within participants to obtain a representative classification accuracy for each participant. Finally, these representative values were averaged across participants. The statistical analysis was performed using sample size of 11, which was equal to the number of observers. To address the multiple comparisons, we adopted the Benjamini-Hochberg (BH) false discovery rate (FDR)-correction method (Benjamini & Hochberg, 1995).

**Figure 4.**
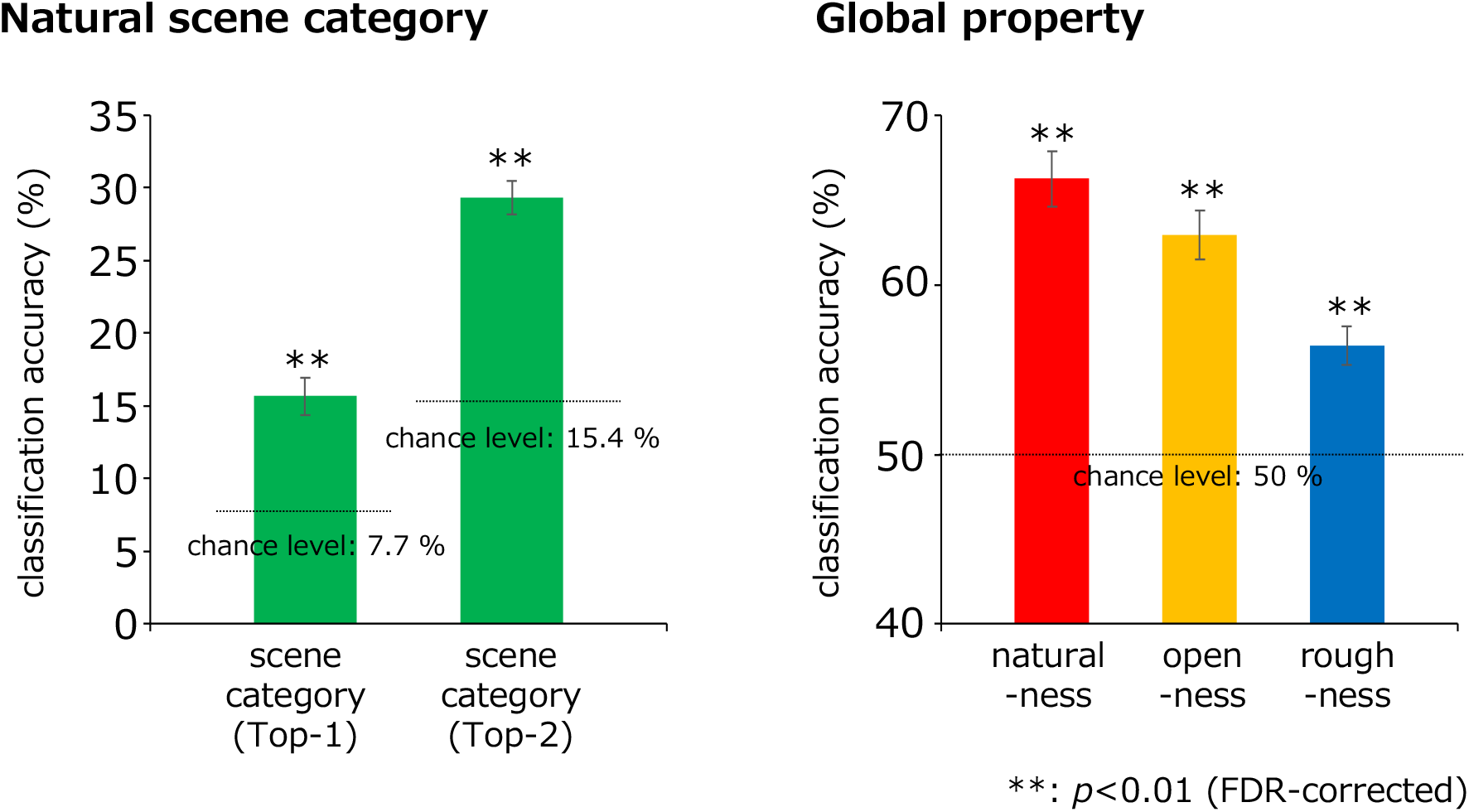
Classification results obtained using the EEGNet models. The green, red, yellow, and blue bars indicate the classification accuracies for the natural scene categories, naturalness, openness, and roughness, respectively, all of which were statistically significant (**: p < 0.01, FDR-corrected). In the classification of the natural scene categories, the Top-n classification accuracy is the rate that the correct category was included in the top-n prediction.

The classification accuracy for the 13 natural scene categories was 15.7 % (chance level: 1/13 (7.7%); p = 1.3×10-4, two-tailed one-sample t-test) and that within the top two categories was 29.3 % (chance level: (2/13 (15.4%); p = 4.7×10-7, two-tailed one-sample t-test), with both results being statistically significant (p < 0.01, FDR-corrected).

Meanwhile, because the train/test split was performed as equally as possible within the natural scene categories, a 0/1 balance in the testing data was not ensured under the naturalness, openness, and roughness conditions. Therefore, to fairly examine the accuracy of the models that classified these global properties, the balanced accuracy calculated using equation (1) was adopted to calculate the classification accuracy for those conditions. Note that tp, fn, fp, and tn denote the numbers of true positives, false negatives, false positives, and true negatives, respectively.

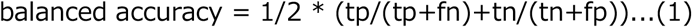

The classification accuracies of naturalness, openness, and roughness were 66.2, 62.9, and 56.4 %, respectively (chance level: 1/2 (50 %); p = 2.2×10^−6^, 6.7×10^−6^, 3.5×10^−4^ respectively, two-tailed one-sample t-test), all of which were statistically significant (p < 0.01, FDR-corrected). These results indicate that natural scene categories and global properties can be significantly classified using simple VEPs.

### Spatiotemporal maps of VEP components contributing to the classification based on Grad-CAM

Figure 5(a) shows the topography of the EEG channels and time points that contributed to the classification visualized using Grad-CAM. The degree of contribution is converted to relative values according to the minimum and maximum values, and we set the minimum values as zero to plot. Red indicates the maximum contribution to the classification, and blue indicates the minimum. Fp1 and Fp2, which were excluded from the classification, are plotted as making the minimum contributions to the classification. In Figure 5(b), the data from the occipital (O1, O2), parietal (P7, P8), and frontal (F3, F4, F7, F8, Fz, Cz) lobes are graphically presented. The data in each panel were averaged across electrodes for visualization.

**Figure 5.**
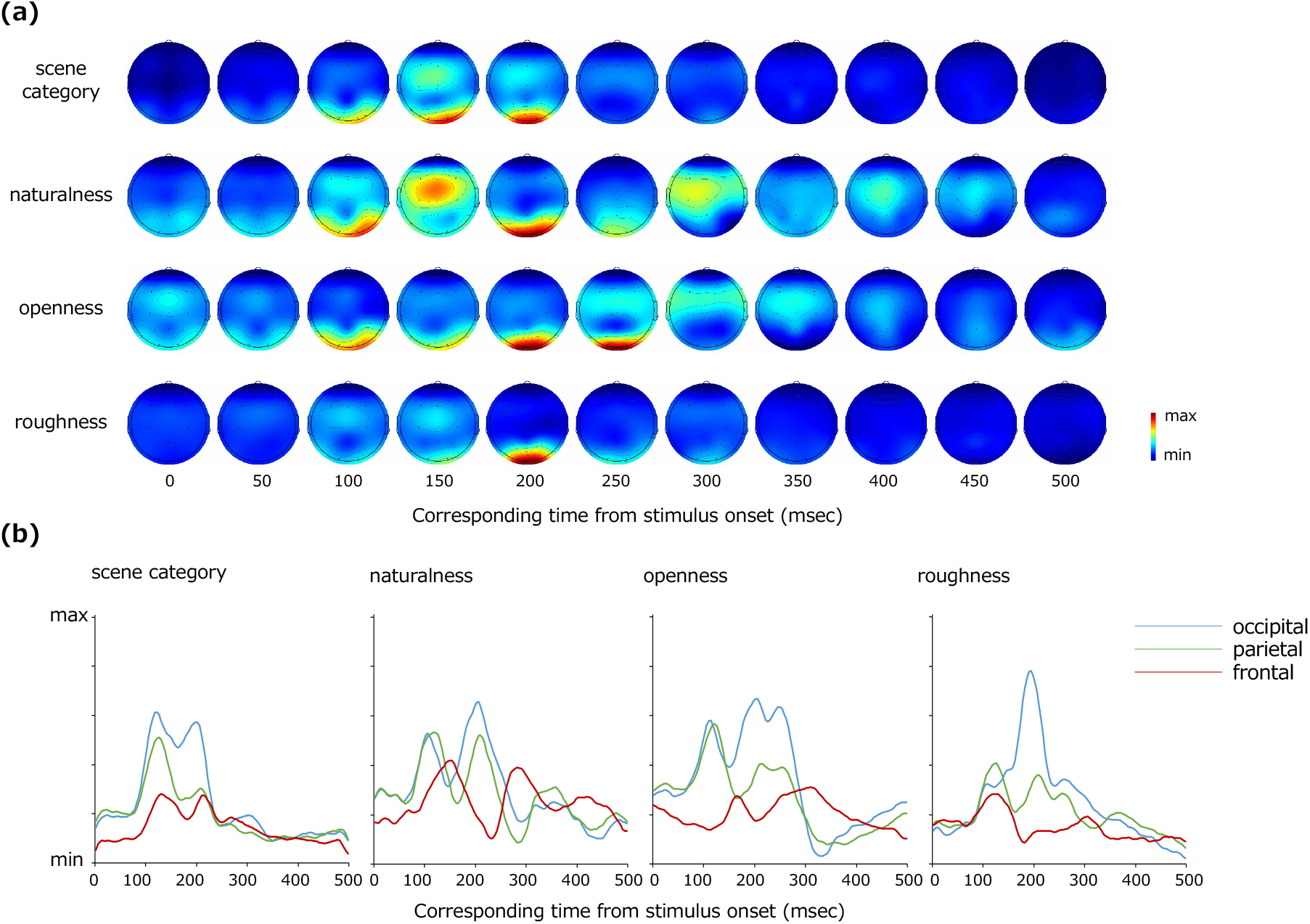
(a) Topography of VEPs contributing to the classification visualized by Grad-CAM. From the top, the EEG channels that contributed to the classification of the natural scene categories, naturalness, openness, and roughness are shown. (b) Re-plot of (a) on a plane with the vertical axis representing the degree of contribution and the horizontal axis representing the time from the stimulus onset. The data were averaged for the occipital (O1, O2), parietal (P7, P8), and frontal (F3, F4, F7, F8, Fz, Cz) lobe channels, respectively.

The occipital and parietal electrodes (O1, O2, P7, P8) at approximately 84–228 ms after the stimulus onset and the frontoparietal electrodes (F3, F4, F7, F8, Fz, Cz) at approximately 124–164 and 200–236 ms contributed to the classification of the natural scene categories.

The frontoparietal electrodes (F3, F4, F7, F8, Fz, Cz) at 144–164 ms after the stimulus onset contributed to the classification of naturalness, and the occipitoparietal electrodes (O1, O2, P7, P8) at earlier latencies such as 92–132 ms after the stimulus onset and relatively late latencies of 172–244 ms also contributed to the naturalness classification. The occipitoparietal electrodes (O1, O2, P7, P8) at approximately 80–284 ms after the stimulus onset contributed to the classification of openness. Furthermore, occipitoparietal electrodes (P7, P8) at approximately 100–132 ms and occipital electrodes (O1, O2) at approximately 164–228 ms after the stimulus onset contributed to the classification of roughness, whereas the other channels did not largely contribute.

These results indicate that the occipital lobe contributed to the classification generally. However, there were certain differences according to the classification classes. As examples, the frontal lobe contributed to the naturalness classification, and the occipital lobe at earlier latencies (approximately 92∼ ms) contributed to the naturalness and openness classification.

## Discussion

To investigate the spatiotemporal development of natural scene perception in the human brain, the present study introduced a deep classification model (EEGNet) that classified natural scene categories and global properties by inputting VEPs for natural scene images. As a result, we found that natural scene categories and global properties can be classified at a statistically significant level even using VEPs with low spatial resolution. We also found that the time points and EEG channels that contributed to the classification differed largely depending on the classes of classification. For example, for naturalness and openness, VEPs in the occipital lobe at early latencies (approximately 80∼ ms) contributed to the classification, whereas VEPs in the occipital lobe at relatively late latencies (∼ 150 ms) mainly contributed to the classification of natural scene categories. In addition, VEPs in the frontal and parietal lobes with late latencies (approximately 164∼ ms) contributed to the classification of naturalness, whereas VEPs in the occipital lobe mainly contributed to the classification of the other classes. These results suggest that different global properties of natural scenes are processed at different latencies and in different areas of the human brain.

In terms of naturalness, the occipital VEPs at earlier latencies (approximately 92∼ ms) and the frontal VEPs at later latencies (approximately 144∼ ms) contributed to the classification. This is consistent with the psychophysical finding that the naturalness of natural scenes can be perceived even with a particularly short presentation (Joubert et al., 2007; Greene & Oliva, 2009; Loschky and Larson, 2010). The reason why naturalness is encoded at early latency may be that naturalness can be predicted to some extent by lower-order image features. In fact, another analysis showed that naturalness was predicted with 85.2% accuracy (chance level: 50%; p = 0.00033, two-tailed one-sample t-test) by the spatial envelope (Oliva & Torralba, 2001; Torralba & Oliva, 2003) using a support vector machine (SVM), which corresponds to the energy of subbands in spatial blocks.

Furthermore, the fact that the VEPs in the frontal lobe contributed the most to naturalness classification supports the results of previous studies using fMRI, which suggested that the parietal and occipital lobes are related to scene recognition by using low-spatial-frequency information (Peyrin et al., 2004, 2010). Man-made scenes contain many linear contours, whereas natural scenes do not necessarily do so. These differences can also be observed even for low-spatial-frequency information, and considering the findings of the previous studies, it is reasonable that the VEPs in the frontal lobe at approximately 150 ms contributed to the naturalness classification.

The openness classification was supported by the occipital VEPs at early latencies (approximately 80∼ ms), which is consistent with the previous finding that natural scene openness modulates the P1 component of the ERP (Hansen et al., 2018). The early encoding of openness may relate to the fact that natural scene openness can be discriminated by lower-order features. In fact, the openness of the visual stimuli was classified with an accuracy of 78.5% (chance level: 50%; p = 0.0011, two-tailed one-sample t-test) by the SD (seven scales, one-octave step) of the spatial frequency subbands of each image using an SVM. The subband SD corresponds to a subset of image statistics known to be important in texture perception in the early visual cortex (Bergen & Adelson, 1988; Heeger & Bergen, 1995; Zipser et al., 1996; Portilla & Simoncelli, 2000; Baker & Mareschal, 2001; Motoyoshi et al., 2007; Freeman & Simoncelli, 2011; Freeman et al., 2013; Ziemba et al., 2019). This image statistic has also been revealed to be strongly correlated with VEPs at as early as 88∼ ms after the stimulus onset (Orima & Motoyoshi, 2021).

Meanwhile, the occipital lobe VEPs at relatively late latency (mainly 164∼ ms) contributed to the classification of roughness, which supports the idea that the roughness is encoded slowly because it is correlated with higher-order information compared with naturalness and openness. Our analysis showed that the roughness of natural scenes was classified with an accuracy of 61.6 % (chance level: 50%; p = 0.0092, two-tailed one-sample t-test) by the cross-spatial-frequency correlation of energy subbands of each image using an SVM. Cross-subband correlation is known as a higher-order image statistic, mainly encoded in V2 (Freeman et al., 2013; Ziemba et al., 2019), and it has been revealed to be strongly correlated with VEPs at later latency (150∼ ms) (Orima & Motoyoshi, 2021). The results in the present study were consistent with these findings.

The classification of the 13 individual natural scene categories was supported by the VEPs of the occipitoparietal electrodes at approximately 84–228 ms, and the frontal lobe at approximately 124–164 and 200–236 ms, which may correspond to the ultra-rapid recognition of natural scenes discussed in the previous studies (Thorpe et al., 1996; Fabre-Thorpe et al., 2001; VanRullen & Thorpe, 2001). These results were mostly a combination of the naturalness, openness, and roughness results and suggested that the natural scene categories reflected the differences in the global properties of natural scenes in a composite manner. The contribution of the occipital lobe to the classification of natural scene categories further supports the results of previous studies that revealed the encoding process of natural scenes by VEPs (Scholte et al., 2009; Hansen et al., 2011; Groen et al., 2013; Greene & Hansen, 2020) and is consistent with the idea that scene selective regions such as the occipital place area, parahippocampal place area, and retrosplenial complex are distributed in the occipital and parietal lobes (Epstein & Kanwisher, 1998; Aguirre et al; Ishai et al., 1999; O’Craven & Kanwisher, 2000; Julian et al., 2016). In addition, the frontal lobe has been suggested to be associated with natural scene perception, which is consistent with the results of previous studies using fMRI (Peyrin et al., 2004, 2010; Walther et al., 2009). The results also suggest that it is possible that natural scenes are processed inter-regionally, from the occipital to frontal lobes.

In the present study, we used RGB images of natural scenes without any control of low-level image measurements across stimuli (e.g., mean luminance and color). Therefore, one may point to the possibility that EEG signals depended on these low-order image features rather than information indicative of the scene category per se. However, such lower-order image statistics are also regarded as a critical attribute of a particular natural scene; e.g., the scenery that most people recall as a coast would be a seaside area in the daytime, which is mostly bright. In this respect, strictly normalizing image features across visual stimuli could rather result in an ecologically invalid finding. Meanwhile, it is noted that our data based on a large but still limited number of images do not completely rule out the possibility that the results were biased by the distribution of lower-order statistics in the image set that we used. In future studies, it may be desirable to obtain more data using a larger image set.

Both psychophysical and physiological studies on scene perception, including the present study, basically use visual stimuli of small size displayed on a conventional computer monitor (e.g., Hansen et al., 2011; Groen et al., 2013). However, given that a goal of scene perception research is to explain our natural scene perception in daily lives, one should ideally use visual stimuli with a sufficiently wide field of view to immerse observers in the scene and allow a high mobility of the observers. It would be difficult to apply such an experimental setting to fMRI experiments that require observers to view stimuli of a limited viewing angle with the head rigidly fixed. In contrast, it may be easier to establish such a free viewing condition with EEG. We expect that the decoding techniques introduced in the present study will also be useful in revealing the cortical dynamics of scene processing in such a natural situation.

